# Efficient GFP-labeling and analysis of spermatogenic cells using the IRG transgene and flow cytometry

**DOI:** 10.1101/465286

**Authors:** Leah L. Zagore, Cydni C. Akesson, Donny D. Licatalosi

## Abstract

Spermatogenesis is a highly ordered developmental program that produces haploid male germ cells. The study of male germ cell development in the mouse has provided unique perspectives into the molecular mechanisms that control cell development and differentiation in mammals, including tissue-specific gene regulatory programs. An intrinsic challenge in spermatogenesis research is the heterogeneity of germ and somatic cell types present in the testis. Techniques to separate and isolate distinct mouse spermatogenic cell types have great potential to shed light on molecular mechanisms controlling mammalian cell development, while also providing new insights into cellular events important for human reproductive health. Here, we detail a versatile strategy that combines Cre-lox technology to fluorescently label germ cells, with flow cytometry to discriminate and isolate germ cells in different stages of development for cellular and molecular analyses.

## Introduction

Spermatogenesis is the development of haploid male germ cells in the testis. Owing to the large number of highly ordered and distinct steps in this developmental program, mouse spermatogenesis has proven to be a powerful model system to study different aspects of mammalian cellular and molecular biology (Youds & Boulton, 2011; Lie, Mruk, Lee, & Cheng, 2010; Song & Wilkinson, 2014; Licatalosi, 2016). This developmental program involves numerous cell divisions and differentiation events driven by intrinsic and extrinsic factors. In mice, germ cells are first specified around day 7 of embryogenesis (E7). These cells, known as primordial germ cells (PGCs), are unique as they are sexually bipotent: capable of differentiating into either oocytes or sperm. Gene expression in somatic cells directs germline sex determination. In a male somatic environment, expression of the SRY gene leads to a male program of germ cell development in which PGCs develop into gonocytes (Kocer, Reichmann, Best, & Adams, 2009; Murray, Yang, & Van Doren, 2010). After a few mitotic divisions, gonocytes become quiescent for the remainder of embryogenesis. A few days after birth, mitosis resumes, gonocytes migrate to the basal membrane of the seminiferous tubule, and a spermatogonial stem cell pool is established that provides the foundation for continued fertility throughout the reproductive life of an organism (Manku & Culty, 2015).

In the postnatal testis, the spermatogenic program can be divided into three general cell stages: mitosis (spermatogonia), meiosis (spermatocytes), and post-meiotic differentiation (spermatids). Spermatogonial stem cells either self-renew to maintain the stem cell pool or divide into differentiating spermatogonia (Busada & Geyer, 2016). Notably, differentiation into A1 spermatogonial cells marks the beginning of the synchronized, time-regulated stages of spermatogenesis. These immature cells progress through a series of mitotic divisions with incomplete cytokinesis to generate chains of A2, A3, A4, Intermediate, and type B spermatogonia. Around 9 days after birth (p9), type B spermatogonial cells undergo one final mitotic division to give rise to primary spermatocytes that enter the second stage of spermatogenesis: meiosis (de Rooij & Griswold, 2012; Griswold, 2016). During meiosis, a single round of DNA replication occurs in spermatocytes, followed by homologous recombination and two successive divisions to produce genetically distinct, haploid round spermatids. Thereafter, spermatids undergo a differentiation program called spermiogenesis, which requires ~13.5 days to complete in mice. During this stage, nuclear repackaging and compaction occur, as well as vast morphological and cytological changes to produce spermatozoa. These include cell elongation, formation of a flagella, redistribution of organelles, and shedding of a large proportion of the spermatid cytoplasm in the form of a cytoplasmic droplet (also called residual body). The cytoplasmic droplet is engulfed by Sertoli cells - somatic cells that span the width of the seminiferous epithelium. Sertoli cells are critical at all stages of spermatogenesis, and provide nutritional and structural support to germ cells as they translocate towards the lumen (Griswold, 1998). Upon completion of spermiogenesis, germ-Sertoli cell contacts are severed to release spermatozoa into the tubule lumen (Upadhyay, Kumar, Ganeshan, & Balasinor, 2012).

Studies of mouse spermatogenic cells have provided key insights into the molecular mechanisms balancing cell proliferation, differentiation, and apoptosis; DNA repair and recombination; cell-cell communication; adhesion; and stage-specific transcriptional and post-transcriptional control of gene expression. For example, molecular studies of round and elongating spermatids have revealed post-transcriptional pathways that temporally control translation of select mRNAs in later stages of spermiogenesis when transcription is silenced (Braun, 1998; Monesi, Geremia, D’Agostino, & Boitani, 1978). Continued study of spermatogenic cells has the promise to provide important insights into many aspects of cell biology, infertility, and testicular cancer (Matzuk & Lamb, 2008; Lin & Matzuk, 2014).

Investigation of cell type specific regulatory programs in heterogeneous tissue requires the ability to distinguish between the numerous cell types present. Several methods exist to isolate and enrich for specific germ cell populations (Meistrich, 1977). STA-PUT, which requires expertise in the morphological identification of germ cells, takes advantage of differences in cell size and sedimentation velocity to separate enriched populations of germ cells with high yields (Bryant, Meyer-Ficca, Dang, Berger, & Meyer, 2013). Immunomagnetic isolation can provide highly pure, viable cell populations, however the number of specific cell types that can be collected is limited (van der Wee, Johnson, Dirami, Dym, & Hofmann, 2001). Further resolution and simultaneous collection of multiple germ cell populations can be achieved using Hoechst-based flow cytometry approaches (Bastos et al., 2005; Gaysinskaya, Soh, van der Heijden, & Bortvin, 2014). For example, cell suspensions from seminiferous tubules can be stained with the DNA-binding dye bis-benzamide Hoechst 33342 (Ho) followed by fluorescence activated cell sorting (FACS) to discriminate, quantify, and collect cells based on DNA content and chromosome structure. The latter is possible due to a chromatic shift in Ho emission (from red to blue) as the concentration of DNA-bound Ho increases (Petersen, Ibrahim, Diercks, & van den Engh, 2004). This allows further discrimination of subpopulations of germ cells with the same DNA content such as primary spermatocytes in different steps of meiotic prophase I, and spermatogonia from secondary spermatocytes (both diploid). A limitation however, is that the emission spectra of diploid germ cells overlaps with that of somatic cells in seminiferous tubules. We have demonstrated that this barrier can be overcome by combining Ho staining with GFP-labeling of germ cells (discussed below).

Cre-lox tools have revolutionized developmental, cellular, and molecular biology research. The general approach uses a single enzyme derived from bacteriophage P1, Cre recombinase, to control site-specific recombination at short sequences (LoxP elements) present in a target gene (Sauer & Henderson, 1988). Depending on the orientation of the LoxP sequences, Cre recombination results in excision or inversion of intervening DNA, thus altering the primary sequence of an endogenous gene or integrated cDNA transgene. Using inducible or tissue-restricted promoters, Cre expression itself can be tightly controlled in a temporal and cell-type specific manner. These approaches are particularly useful when whole-animal gene deletion results in embryonic or perinatal lethality, thus preventing analysis of the roles of these genes in postnatal steps of spermatogenesis. In addition to ablating expression of specific genes, Cre-lox can be used to turn on expression of transgenes or create epitope-tagged proteins. For example, in Brainbow mice Cre-lox allows hundreds of individual neurons to simultaneously have a distinct fluorescence signature to efficiently map neural circuitry in the brain (Livet et al., 2007). These strategies have also allowed for detailed study into cell type specific alternative gene regulation and processing using intact, heterogeneous tissue. For example, cTAG-PAPERCLIP uses Cre-lox to conditionally tag polyA binding protein (PABP) with GFP to profile alternative mRNA species in select populations of cells (Hwang et al., 2017).

Popular Cre-drivers in spermatogenesis research include those that restrict Cre expression to Sertoli cells (Amh-Cre (Lécureuil, Fontaine, Crepieux, & Guillou, 2002)) or germ cells at different developmental time points (such as Stra8-iCre (Sadate-Ngatchou, Payne, Dearth, & Braun, 2008), Ddx4-Cre (Gallardo, Shirley, John, & Castrillon, 2007), Prdm1-Cre (Ohinata et al., 2005) and Prm1-Cre (O’Gorman, Dagenais, Qian, & Marchuk, 1997). We previously used the Stra8-iCre driver and the IRG transgene (a LoxP-containing dual fluorescence reporter that switches expression from red to green fluorescent protein after Cre-recombination) (De Gasperi et al., 2008) to label postnatal germ cells with GFP. We first used this strategy with Ho staining and FACS, to quantify subpopulations of germ cells from adult mice with germ cell-specific deletion of *Ptbp2*, which encodes an essential RNA binding protein (Zagore et al., 2015). More recently, we used the Stra8-iCre and IRG transgenes to GFP-label and collect the limiting numbers of spermatogonia present in p6 testes of *Dazl* knockout mice (Zagore et al., 2018). In this report, we expanded our analysis of GFP expression in the postnatal testis of Stra8-iCre+, IRG+ mice to further assess the utility of IRG-based GFP-labeling as a tool for spermatogenesis studies. In addition to showing that GFP is present in germ cells in each of the main stages, we show that GFP intensity markedly increases during spermatid elongation. Furthermore, we demonstrate that the difference in GFP intensity between round and elongating spermatids provides an additional FACS parameter that can be used to quantify and isolate round versus elongating populations of spermatids. Lastly, we show that the IRG transgene can be also be used in combination with the Ddx4-Cre driver to label gonocytes during embryogenesis. Collectively, our findings demonstrate that GFP-labeling of germ cells with the IRG transgene and germ cell-specific Cre drivers is an effective approach to discriminate, quantify, and collect germ cells at different stages of development.

## Results

### GFP-labeling mitotic, meiotic and post-meiotic germ cells with Stra8-iCre and IRG

To assess GFP expression during the first wave of spermatogenesis, we used flow cytometry to examine Ho-stained preparations of cells from Stra8-iCre+ IRG+ seminiferous tubules collected at p23, p35, and p42 (Figure 1). After gating for GFP+ cells, three diagonal bands of cells corresponding to 1C, 2C, and 4C cells are evident, as expected (Zagore et al., 2015). Whereas the 1C band contains a single cluster of GFP+ cells, each of the 2C and 4C cells segregate into sub-clusters with either low or high Ho-red fluorescence emission. The five readily discernible cell clusters correspond to: 1) spermatogonia (2C, mitotic), 2) pre-leptotene/leptotene spermatocytes (4C, early meiosis I prophase), 3) pachytene/diplotene spermatocytes (4C, early meiosis I prophase), 4) secondary spermatocytes (4C, late meiosis I prophase), and 5) spermatids (1C, post-meiotic) (Figure 1A). Importantly, the percentage of germ cells in each of the five subgroups differed between testes collected at different ages and closely paralleled reported values (Bellvé, Millette, Bhatnagar, & O’Brien, 1977; Bellve et al., 1977) (Figure 1B-E). Consistent with the fact that a minority of germ cells have completed meiosis by p23, only 15.7% of the GFP+ cells were present in population 5 corresponding to haploid spermatids (Figure 1B). In contrast, haploid cells comprised a significantly greater proportion of germ cells present in testes from p42 mice, consistent with completion of the first wave of spermatogenesis at ~p35 (Figure 1C). Another significant shift was readily observable in the development of early prophase-I spermatocytes, which decreased from approximately 25% to less than 8% of GFP+ cells at p42. Importantly, the relative levels of GFP+ germ cells in different stages of spermatogenesis identified by Ho-staining and FACS are entirely consistent with relative germ cell numbers reported by cytological analysis of germ cells present in mouse testes at different developmental time points (Bellvé et al., 1977). These observations show that GFP is continually present throughout spermatogenesis following recombination of the IRG transgene in spermatogonia. They also indicate that dual fluorescence FACS analysis of Stra8-iCre, IRG cells is an effective strategy to discriminate and quantify subpopulations of germ cells in different stages of spermatogenesis.

**Figure 1.**
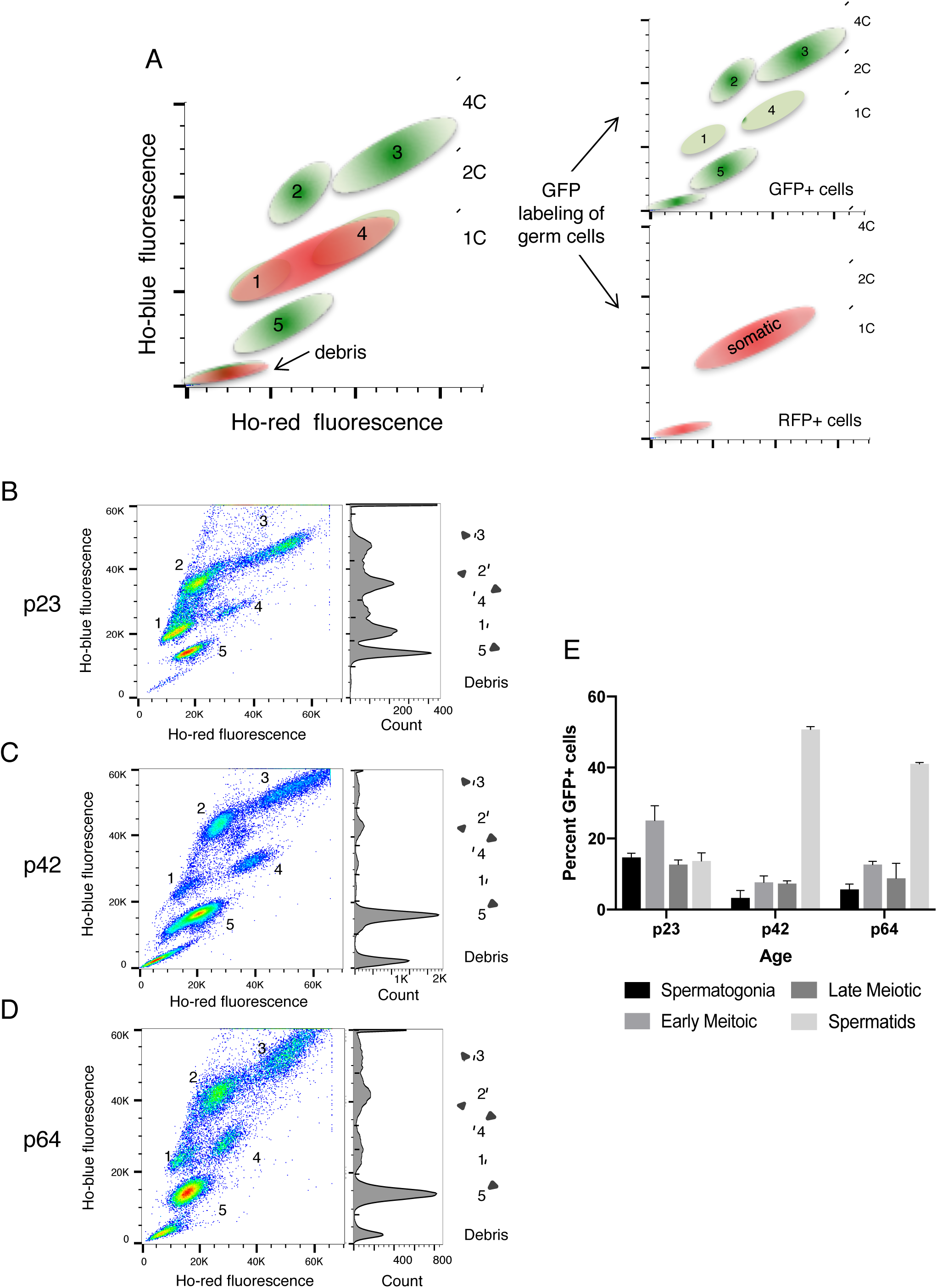
Quantitative analysis of the FACS profiles from juvenile and adult testes using the dual fluorescence reporter system. **A.** Representative image showing a FACS profile from Hoechst 33342 stained adult male testis lysate. Cells cluster into 5 subpopulations: 1) spermatogonia, 2) early-prophase I spermatocytes, 3) late-prophase I spermatocytes, 4) secondary spermatocytes, and 5) spermatids. Note the resolution of the GFP+ diploid population using fluorescence labeling of germ cells. **B, C, D.** Distribution of GFP positive, Hoechst 33342-stained cells from C57 Stra8-iCre+, IRG+ mice at ages p23, p42, and p64 respectively. Histogram to the right shows the distribution of GFP+ cells based on Ho-blue fluorescence values. **E.** Percentage of total GFP positive cells within specific gated germ cell populations. Percentage reported reflects the average percentage of two biological replicates and error bars represent standard deviation.

We next used fluorescence microscopy to examine GFP-expression in Stra8-iCre+, IRG+ testes collected at postnatal days 6, 11, 19, 29, and 35 (Figure 2). At p6, type A spermatogonia are the most advanced germ cells present in seminiferous tubules. By p11, meiotic prophase has initiated, and by p19 germ cells have reached the late pachytene stage. Round and elongating spermatids are readily observable at p29. Finally, p35 marks the completion of the first wave of spermatogenic cell development. Sections were incubated with antibody to GFP and counterstained with antibody for the germ cell marker DDX4. Consistent with Stra8-iCre expression beginning ~p3 (Sadate-Ngatchou et al., 2008), GFP expressed from the IRG transgene was observed in spermatogonia present in p6 testes. Importantly, GFP expression was restricted to DDX4+ cells, and greater than 95% of DDX4+ cells were also positive for GFP across multiple ages and biological replicates. (Table 1) The presence of a small subset of DDX4+, GFP-cells is consistent with previous reports that Stra8-iCre is not expressed in a subset of undifferentiated spermatogonia (Sadate-Ngatchou et al., 2008). As expected, the number of GFP+ cells per tubule expanded with increased age as germ cells advanced through spermatogenesis. Therefore, in agreement with FACS analysis, GFP was observed in all stages of postnatal germ cell development.

**Figure 2.**
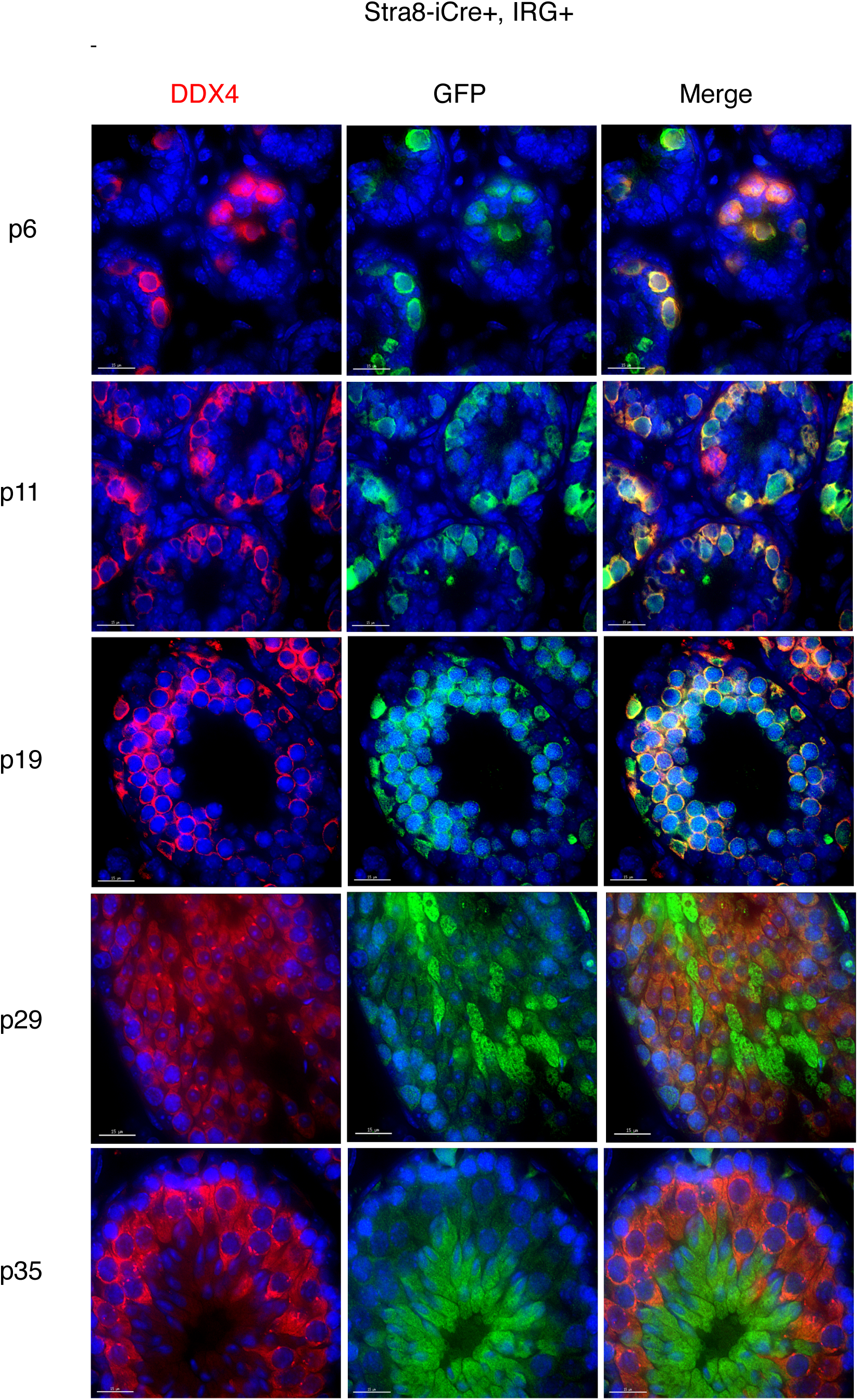
Stra8-iCre mediated GFP expression throughout the first wave of spermatogenesis in Stra8-iCre, IRG+ mice. **A-E.** Representative images of GFP and Ddx4 expression in seminiferous tubule crosses sections at ages p6, p11, p19, p29, and p35 are shown. Note that because of a large dynamic range in GFP expression, GFP intensity is not to scale between different age time points. Scale bars represent 15 μm.

**Table 1.** Quantification of Cre-mediated recombination of the IRG transgene in germ cells. Germ cells were identified as DDX4+ cells and the overlap between GFP+ cells and DDX4+ cells was calculated using two biological replicates. The high number of cells both GFP+ and DDX4+ provide a measure of the efficiency recombination and cell restricted Cre-expression.

| Cre Driver | GFP+, DDX4+ | GFP-, DDX4+ | GFP+, DDX4- | # cells counted | # tubules counted | ages |
| --- | --- | --- | --- | --- | --- | --- |
| Stra8-iCre | 97.32% | 2.5% | 0.180% | 560 | 39 | p7,p11,p19 |
| Ddx4-Cre | 94% | 1.71% | 4.27% | 117 | 37 | p0 |

### Differential GFP levels mark round and elongating spermatids

Immunofluorescence microscopy shows that GFP intensity was comparable in the cytoplasm of spermatogonia, spermatocytes, and round spermatids. Unexpectedly however, GFP intensity showed a pronounced increase in elongating spermatids. A key step in the development of spermatozoa is proper formation of the acrosome – a Golgi and endosome derived vesicle which first appears as a pro-acrosomal granule, then attaches and extends over the nuclear membrane. We used fluorophore-conjugated PNA to classify spermatids in each of the 16 defined steps of differentiation and determine the precise step when GFP levels increase (Nakata, Wakayama, Takai, & Iseki, 2015; Meistrich & Hess, 2013)(Figure 3A-E). We consistently found that increased GFP intensity was first observed in spermatids at step 8 (Figure 3D), when the acrosome has spread over more than 1/3 over the nucleus and some round spermatids have oriented toward the basal membrane. Thereafter, GFP intensity continually increased and in step 10 elongating spermatids show a marked difference compared with pachytene spermatocytes (Figure 3E).

**Figure 3.**
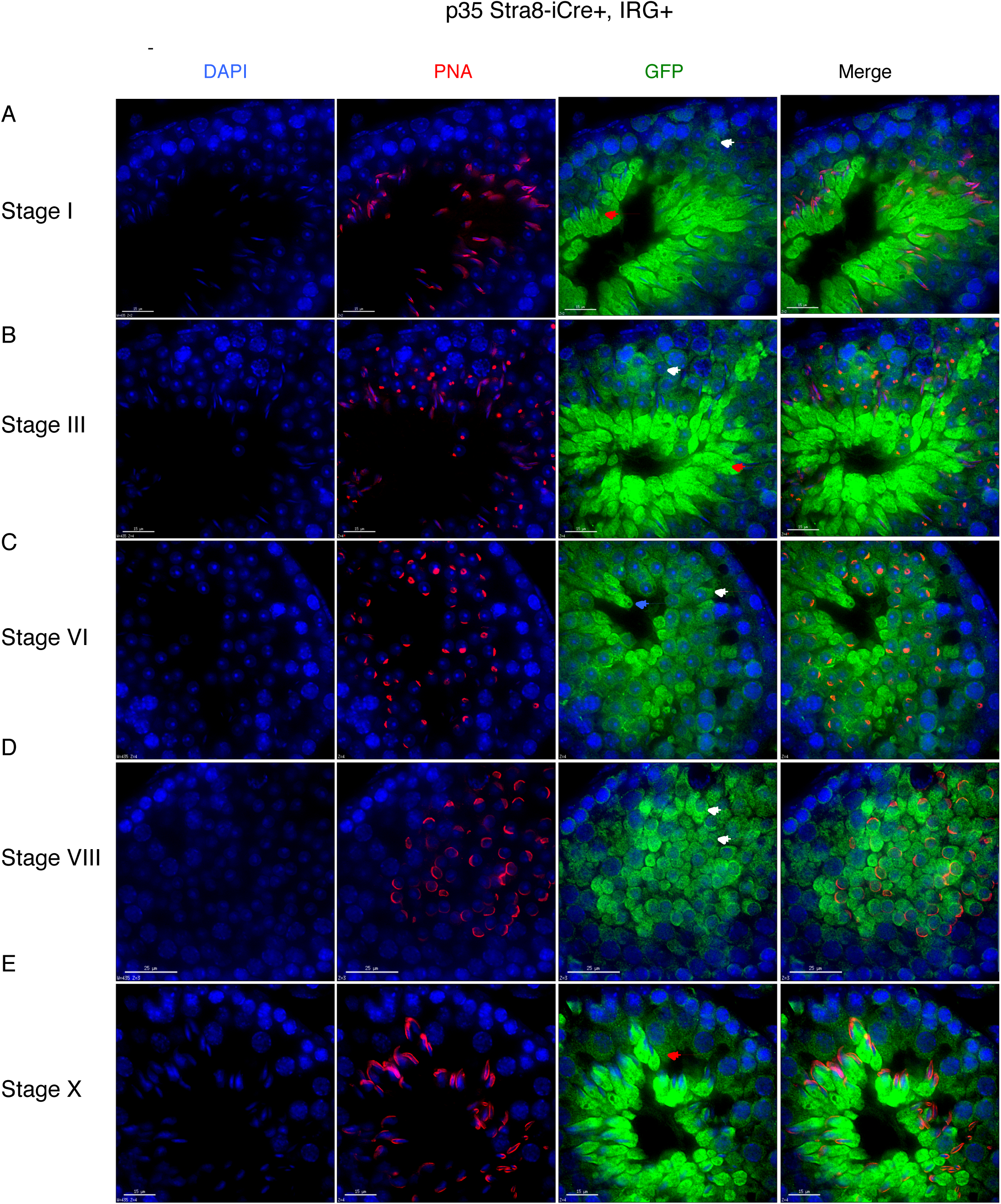
Staging of seminiferous epithelial cells to pinpoint the timing of increased GFP expression. Cross-sections from p35 Stra8-iCre+, IRG+ mouse testes were stained for GFP and peanut agglutinin (PNA) was used to assess acrosomal status. Representative images of spermatogenic stages I, III, VI, VIII, and X are shown **(A-E).** In spermatogenic stages with multiple populations of spermatids, distinct differences in GFP expression can be observed between round spermatids (white arrows – spermatid differentiation steps 1-8) and elongating spermatid (red arrows – spermatid differentiation steps 9-16) populations, with higher GFP signal in the latter **(A,B).** Note the cells with high GFP expression in Stage V represent the residual bodies of Step 15-16 spermatids, as they contain no nucleus (**C**, blue arrow). The subtle difference in GFP expression first begins when round spermatids progress to step VIII **(D,** compare white arrows). Scale bars represent 15 μm.

During spermatid differentiation, most mRNAs assemble into translationally-repressed ribonucleoprotein complexes (RNPs) in a sequence-independent manner (Schmidt, Hanson, & Capecchi, 1999). Only select mRNAs are known to be translated in elongating spermatids. One such example is *Prm1* mRNA, which is made in round spermatids but maintained in a translationally-repressed state, then activated for translation in elongating spermatids. Precocious translation of *Prm1* mRNA results in defects in nuclear repackaging (Lee, Haugen, Clegg, & Braun, 1995). The observed increase in GFP intensity in elongating spermatids was therefore unexpected, and prompted us to ask if GFP mRNA evades the widespread translational repression that occurs in these cells. To test this possibility, we used sucrose density gradient centrifugation to fractionate mRNAs based on the number of associated ribosomes, followed by qRT-PCR. As expected (Kleene, 1989), *Prm1* mRNA sedimented in the RNP fraction in lysates from p23 testes (where the most advanced germ cells present are still in the round spermatid stage), and increased in polyribosome-containing fractions in lysates from p33 testes (which contain elongating spermatids) (Figure 4B). In contrast, GFP mRNA had decreased association with polyribosomes in p33 compared to p23 lysates (from 66% to 22%), indicating decreased translation of GFP protein. Therefore, the increase in GFP intensity is likely due to redistribution and concentration of pre-existing GFP in the cytoplasm of condensing spermatids, rather than increased translation of GFP mRNA.

**Figure 4.**
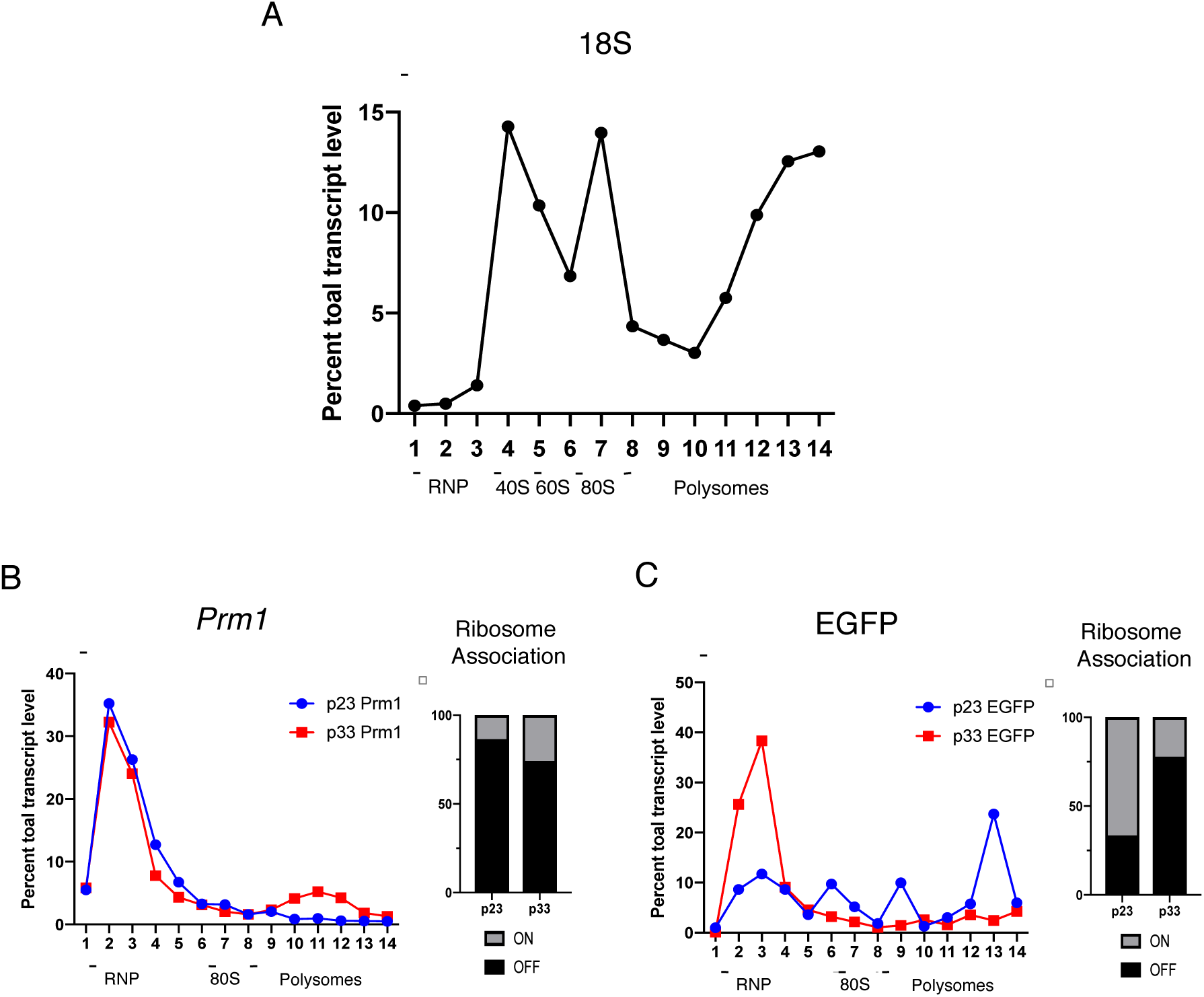
Polysome association of GFP transcripts in round and elongating spermatids. **A.** Distribution of 18S ribosomal RNA across a 10-45% sucrose gradient **B.** Distribution of Prm1 transcripts across sucrose gradient from p23 Stra8-iCre+, IRG+ mouse testes. Right panel shows overall ribosome association of Prm1 at p23 and p33 age males. **C.** Distribution of EGFP transcripts across a 10-45% sucrose gradient from Stra8-iCre+, IRG+ mouse testes. Right panel shows overall ribosome association of Prm1 at p23 and p33 age males. “OFF” ribosome was calculated as the sum of total transcripts from fractions 1-5. “ON” ribosome was calculated as the sum of total transcripts from fractions 6-14. Percentage total transcript level reported is the average of two technical replicates.

Since the fluorescence emission spectra of Ho-stained spermatids yields a single cluster of cells (Figure 1), we asked if the differences in GFP intensity observed in round and elongating spermatids by immunofluorescence could be exploited as a parameter to separate these two sub populations of haploid cells by FACS. To address this question, we re-examined FACS profiles from Stra8-iCre+, IRG+ testes collected from mature and p23 mice, where the majority of spermatids in the latter have not yet progressed to the elongation steps of differentiation (Figure 4). Consistent with immunofluorescence microscopy, spermatids from p23 contained a single population of GFP+ cells (Figure 5A), while spermatids from older mice separated contained a second population of higher GFP-intensity (Figure 5B, C). These observations indicate that differential GFP levels in round and elongating spermatids can be used a parameter to quantify and collect post-meiotic germ cells for downstream cytological and molecular analyses.

**Figure 5.**
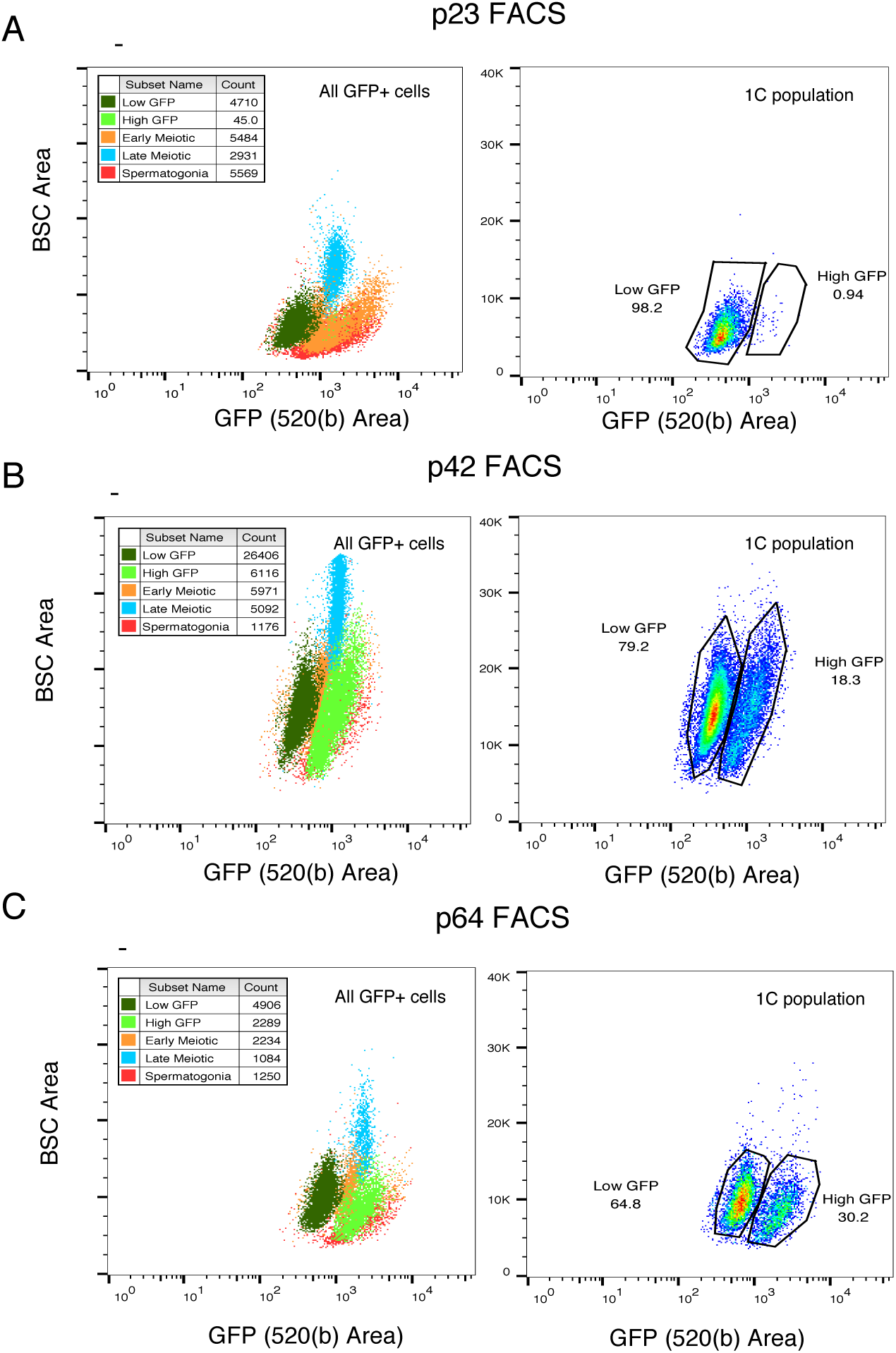
Analysis of GFP intensity in Hoechst 33342-stained GFP+ spermatids. (**A-C**). The left panel shows the GFP intensity of all 5 subpopulations of germ cells, dark green: round spermatids, light green: elongating spermatids, orange: early meiotic spermatocytes, blue: late meiotic spermatocytes, red: spermatogonia in p23, p42, and p64 Stra8-iCre+, IRG+ mice. GFP intensity (520(b) area) of spermatids populations from Figure 1 (B-D, group 5) was assessed. The right panel shows the percentage of spermatids categorized as “Low GFP” or “High GFP” representing round spermatids and elongating spermatids. Note the absence of elongating spermatids (High GFP) at p23.

### Characterization of GFP expression patterns in Ddx4-Cre driver mice

The results described above highlight the utility of the IRG and Stra8-iCre transgenes to GFP-label postnatal germ cells. To determine if the IRG transgene is expressed at earlier time points and might serve as a useful tool to label germ cells during embryogenesis, IRG+ mice were bred with mice containing the Ddx4-Cre transgene, which expresses Cre recombinase around E15.5 (Gallardo et al., 2007). Indeed, p0 gonocytes in Ddx4-Cre+, IRG+ males showed strong, cytoplasmic GFP staining (Figure 6). Notably, the majority of GFP expression at p0 and p6 was restricted to germ cells (94%), indicating efficient recombination and minimal ectopic Cre expression (Table 1). These observations confirm expression of the IRG transgene and GFP-labeling in gonocytes, thus providing a means to identify and collect gonocytes from perinatal mice.

**Figure 6.**
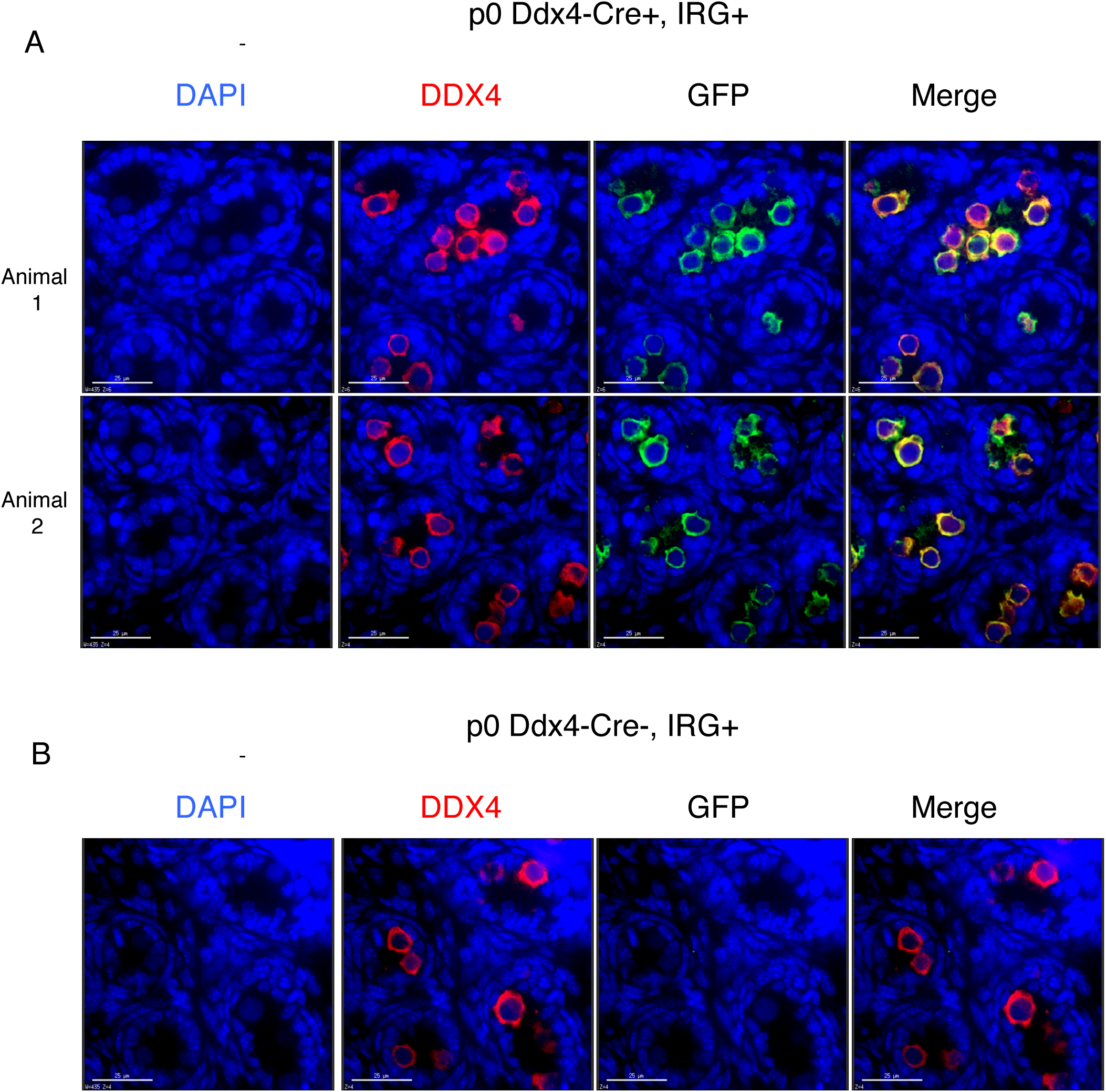
Ddx4-Cre mediated GFP expression in p0 Ddx4-Cre+, IRG+ mice. **A.** Representative images of GFP and DDX4 expression in seminiferous tubule cross-sections at age p0 from two biological replicates. **B.** Negative control showing lack of GFP expression in p0 Ddx4-Cre-, IRG+ mice. Scale bars represent 25 μm.

## Discussion

Site-specific DNA recombination using Cre-lox technology is a powerful method to create knockout or knock-in alleles of endogenous genes. Combining cell type specific Cre drivers with transgenes that express beta-galactosidase or fluorescent proteins has facilitated studies of cell lineages during development and isolation of select cell populations by FACS. These methods are particularly useful for the study of specific cell types in tissues with high cellular heterogeneity. Previously, we demonstrated that FACS analysis of Ho-stained GFP+ cells from Stra8-iCre+, IRG+ testes can uncover subtle, yet quantitative differences in spermatocyte populations between wild type and *Ptbp2* knockout mice not readily discernable by histological approaches (Zagore et al., 2015). In addition, we showed that the Stra8-iCre and IRG transgenes can be used in combination to label and collect limiting numbers of spermatogonia present in *Dazl* knockout mice (Zagore et al., 2018). To further characterize GFP expression from the IRG transgene in germ cells, here we took advantage of the fact that the first round of germ cell development in postnatal mice occurs in a coordinated wave of synchronized and timed events and lasts ~35 days. We show that flow cytometry of Ho-stained GFP+ cells recapitulates known variations in germ cell subpopulations that occur as germ cells progress through the first wave of spermatogenesis. Together, these prior observations and those reported here, confirm the utility of Ho staining of IRG mice and flow cytometry as a quantitative tool for comparative analyses of germ cell development.

While Cre-lox recombination approaches have been implemented in countless studies, expression of Cre does not always follow the expected expression pattern dictated by a given promoter (Smith, 2011). Furthermore, concerns arise as many publications report complications, such as unexpected Cre expression, variable Cre recombination efficiency, and potential toxicity (Schmidt-Supprian & Rajewsky, 2007; Heffner et al., 2012; Matthaei, 2007; Eckardt et al., 2004). Both Stra8-iCre and Ddx4-Cre drivers performed well in terms of recombination efficiency and cell-type specific recombination of the IRG transgene. However, GFP+ DDX4-cells were more prevalent using the Ddx4-Cre driver as compared to Stra8-iCre driver (4.27% vs 0.18%). Consequently, if early postnatal labeling is not required for subsequent analysis, Stra8-iCre may prove a better choice for germ cell restricted Cre recombinase expression. A strong maternal effect has been described for Ddx4-Cre, with prevalent recombination present in some animals (Gallardo et al., 2007); therefore, care should be taken to account for parent-of-origin effects in experimental design. These Cre-drivers could be replaced with other temporally regulated drivers and coupled with FACS; however, further characterization is necessary to assess ectopic Cre expression and unanticipated effects.

We found that differences in GFP levels discriminate round from elongating spermatids, thus providing a simple means to separate these two populations. The timing of increased GFP intensity coincides with the onset of nuclear repackaging, whereby chromatin compaction is driven by the sequential replacement of histones by transition proteins and protamines, resulting in widespread transcriptional inactivation. Synthesis of transition proteins and protamines is dependent on precise, stage-specific dissociation of their mRNAs from mRNPs and association with polyribosomes. While it is known that precise temporal regulation of these mRNAs depends on 3’UTR elements, and is essential for proper chromatin compaction and spermatozoal head formation, the underlying mechanisms remain poorly understood. Exploiting differences in GFP levels in round versus elongating spermatids from Stra8-iCre+, IRG+ mice presents an opportunity to readily separate, quantify, and collect these cells by FACS to further study of post-transcriptional mechanisms during spermiogenesis. Importantly, since spermatids are the only haploid cells in the testis, a chromatic shift in Ho fluorescence emission is not required to distinguish GFP+ spermatids from other germ or somatic cell types, indicating that this approach should be compatible with a broader range of DNA binding dyes and flow cytometers.

## Materials and Methods

### Animals and tissue collection

Mice bearing the *Stra8*-*iCre* transgene (stock Tg[Stra8-icre]1Reb/J0, IRG transgene (B6.Cg-Tg[CAG-DsRed,-EGFP]5Gae/J), and Vasa-Cre transgene (FVB-Tg[Ddx4-cre]1DCas/J) were purchased from The Jackson Laboratory. C57BL6 Stra8-iCre++ males were crossed with C57BL6 IRG+ females to generate C57BL6 Stra8-iCre+, IRG+ offspring. FVB Vasa-Cre++ males were crossed with C57BL6 IRG+ females to generate 50:50 FVB:C57BL6 Vasa-Cre+, IRG+ animals. Mice were sacrificed by isoflurane inhalation followed by cervical dislocation or decapitation. Day of birth was considered p0. All animal procedures were approved by the Institutional Animal Care and Use Committee at CWRU.

### Immunofluorescence microscopy

Testes were detunicated in cold Hank’s balanced salt solution (1× HBSS) and fixed overnight at 4°C in 4% paraformaldehyde. Following fixation, tissue was washed with cold 1X phosphate-buffered saline (PBS) and embedded in paraffin by the Histology Core Facility at CWRU. Immunofluorescence microscopy was performed as previously described (Zagore et al., 2018).

### Flow cytometry analysis of seminiferous epithelial cells

Dual fluorescent FACS approach of Hoechst 33422 (Ho) stained seminiferous tubule cells was performed as previously described (Zagore et al., 2015).

### Sucrose Gradient Polysome Analysis

Mouse testes were collected and detunicated in 1X cold HBSS and flash frozen on dry ice. Tissue was lysed in 320 μL lysis buffer (20 mM HEPES, 100 mM NaCl, 3 mM MgCl_2_, 05.% Triton X-100, 100 μg/mL emetine) and disrupted using a 1 mL syringe plunger. Lysates were incubated on ice for 5 minutes after the addition of 4 μL RNasin (Promega). 200 μL lysate was loaded onto a 10%-45% sucrose gradient and centrifuged for 2 hours at 4oC at 40,000 rpm in a Sw41Ti rotor. After centrifugation the gradient was fractionated into 16 fractions (1^st^ and 16^th^ fractions discarded) and stored at −80°C. 25% of each fraction was used for Ribozol (VWR) RNA isolation followed by isopropanol precipitation. Residual DNA was removed using Turbo DNase (ThermoFisher). qRT-PCR analysis was performed as previously described (Zagore et al., 2015; Zagore et al., 2018) using the following primers: Prm1 F: CGCCGCTCATACACCATAAG, Prm1 R: GGCGAGATGCTCTTGAAGTC, EGFP F: GGTGAACTTCAAGATCCGCC, EGFP R: CTCAGGTGGTTGTCGGG. 18S ribosomal RNA was assessed using a TaqMan Gene Expression Assay designed for mouse Rn18s (ThermoFisher).

## Acknowledgements

We are grateful to the following Case Western Reserve University core facilities: Virology, Next Generation Sequencing and Imaging Core (IF microscopy), Cytometry and Microscopy (FACS), and Tissue Resources (embedding). This work was supported by funds from the NIH to L.L.Z. (T32 GM08056) and D.D.L. (R01 GM107331).

## Author Contributions

L.L.Z and D.D.L conceived of and designed the study. L.L.Z performed the experiments. L.L.Z. and C.C.A performed genotyping and collected tissue for embedding. L.L.Z. and D.D.L wrote the manuscript.

